# The dimensions of species diversity

**DOI:** 10.1101/400481

**Authors:** Matthew J. Larcombe, Gregory J. Jordan, David Bryant, Steven I. Higgins

## Abstract

Diversification processes underpin the patterns of species diversity that fascinate biologists. Two competing hypotheses disagree about the effect of competition on these processes. The bounded hypothesis suggests that species diversity is limited (bounded) by competition between species for finite niche space, while the unbounded hypothesis proposes that evolution and ecological opportunity associated with speciation, render competition unimportant. We use phylogenetically structured niche modelling, to show that processes consistent with both these diversification models have driven species accumulation in conifers. In agreement with the bounded hypothesis, niche competition constrained diversification, and in line with the unbounded hypothesis, niche evolution and partitioning promoted diversification. We then analyse niche traits to show that these diversification enhancing and inhibiting processes can occur simultaneously on different niche dimensions. Together these results suggests a new hypothesis for lineage diversification based on the multi-dimensional nature of ecological niches that accommodates both bounded and unbounded diversification processes.

---

Species diversity has changed dramatically over geological time^1^. Although diversity has clearly increased since life began, reconstructions using the fossil record are ambiguous about the causes of, and constraints on, this increase^2–4^. One important open question is whether the rate of species accumulation slows as diversity increases, or is independent of diversity^4–6^. The latter *unbounded hypothesis* implies that time, and the rate of evolution within clades (monophyletic branches of phylogenies) control diversification and that there is essentially no limit on total diversity^3^. Alternatively, the former *bounded hypothesis* suggests that competitive ecological processes result in a diversity-dependent ceiling on species richness^7^. Resolving this debate is essential for understanding limits to biodiversity, and why diversity is unevenly distributed in space and time and between clades.

Previous attempts to discriminate between bounded and unbounded diversification have focused on modelling species accumulation inferred from phylogenies^8, 9^ and fossil assem-blages^5, 6, 10^, and to a lesser extent testing how ecological niche evolution impacts diversi-fication^11, 12^. The results to date have been inconclusive and often contradictory^2–4^, ^13, 14^, suggesting that a more nuanced explanation may be required^4, 14^. Here we quantify the extent to which both bounded and unbounded processes influence species accumulation in the conifers. Our analysis exploits methodological advances that allow us to infer multi-dimensional physiological-niche properties for large suites of species^15, 16^. We use this data to discriminate between the distinctive niche-characteristics predicted by the bounded and unbounded hypotheses. Specifically we test support for the bounded hypothesis’ prediction that diversification should slow as niche overlap increases within clades^2, 17^ and the unbounded hypothesis’ prediction that niche evolution accommodates increasing diversity by allowing the partitioning or expansion of niche space^3, 18, 19^.

Conifers are an ecologically important, globally distributed division of plants (Fig. 1) that are ideal for this analyses. This large, well-studied lineage has well-defined clades, excellent distribution data^20^, and is ancient enough (> 300 myo;^21^) to assess how species accumulate through time. We use distribution data and a process-based species distribution model (SDM) to infer physiological niche parameters for each of 455 conifer species (75 % of extant conifers). The niche parameters are combined with a robust fossil calibrated phylogeny^21^, and interpreted statistically using an *a-priori* conceptual model of how niche and phylogenetic parameters relate to species richness (Fig. 2). The model postulates that species richness can be impacted both directly and/or indirectly by clade age (CA), the multivariate niche evolution rate (NER), as well as two novel metrics: clade niche size (CNS) and phylogenetic competition index (PCI). Clade niche size is the projected potential niche size (number of grid cells occupied by all species in the clade) corrected for clade species number (see Methods). The phylogenetic competition index is the product of niche overlap and geographic overlap between species within clades. The parameters of this model are estimated using phylogenetically constrained Bayesian path analysis. We conduct the analysis at two phylogenetic levels, using 10 large clades and 42 smaller clades.

**Figure 1.**
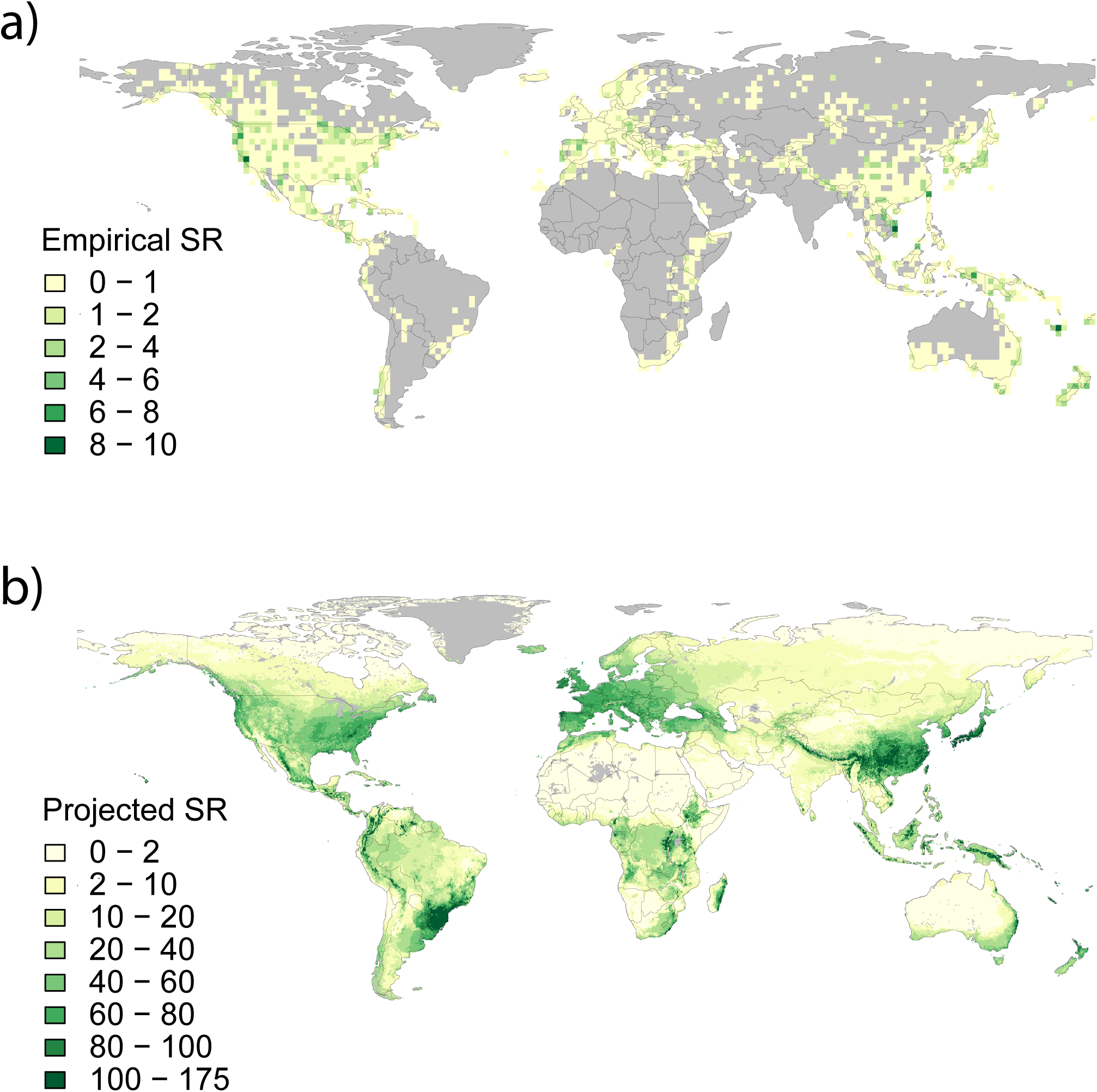
Global species richness (SR) for 455 conifer species based on: a) the cleaned empirical distribution data; and b) projections form process based species distribution models.

**Figure 2.**
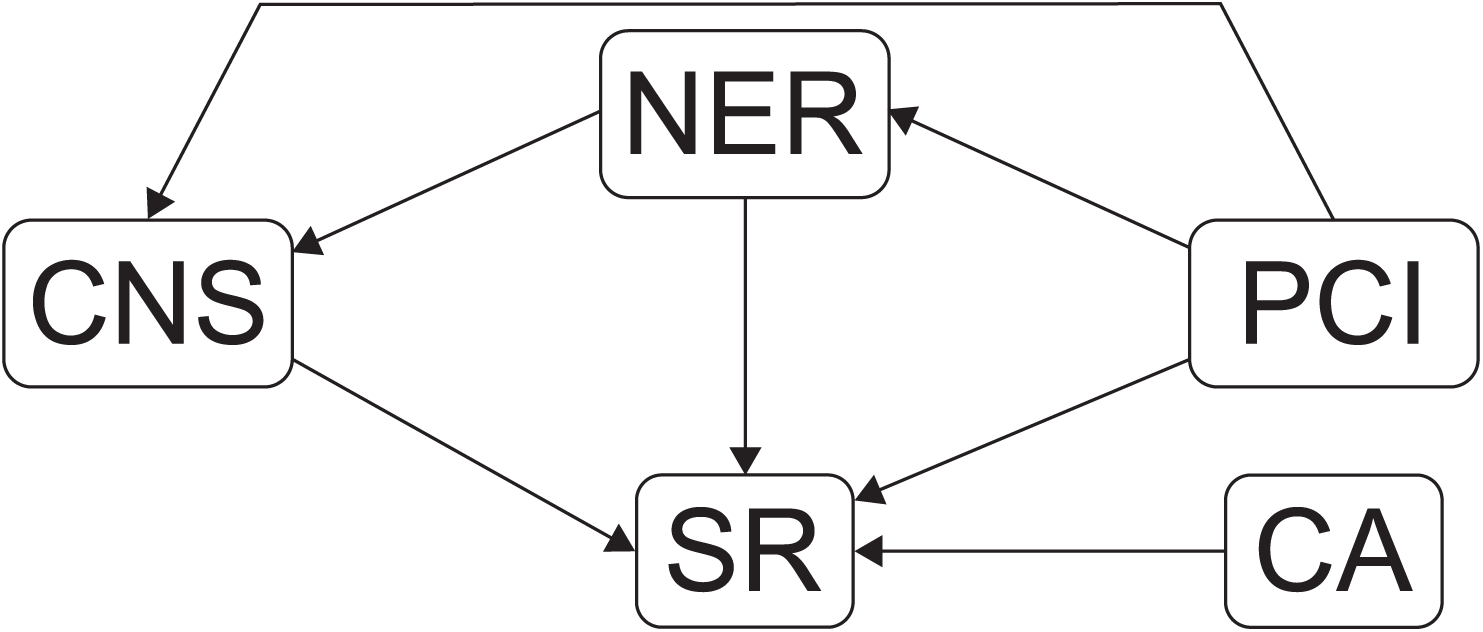
*A-priori* conceptual model of direct and indirect effects of niche and phylogenetic parameters on clade species richness (SR). Bounded diversification predicts a negative effect of PCI, and unbounded diversification a positive effect of NER in combination with variation in CNS, indicating niche partitioning and/or expansion. SR = species richness; CA = clade age; PCI = phylogenetic competition index; NER = multivariate niche evolution rate; CNS = clade niche size.

## Results

We found that diversification in conifers was influenced in almost equal measure by bounded and unbounded processes (Fig. 3a). In line with the bounded hypothesis, competition with relatives (PCI) had a strong negative effect on species richness, which suggests that available niche space can limit species accumulation. This effect was strong in both the 10 (−1.02) and 42 (−0.88) clade analyses. Support for the unbounded hypothesis was evidenced by our finding that niche evolution rate (NER) contributed positively to species richness, suggesting that higher niche evolution rates within clades allow more species to accumulate. This effect was stronger in the 42 clade analysis (0.56) than in the 10 clade analysis (0.31). Furthermore we found that CNS had neutral or negative influence on SR, suggesting that niche partitioning constitutes the main mode of niche evolution in conifers. The negative effect of clade niche size in the 10 clade analysis is somewhat counter-intuitive since it suggests that clades with smaller niche volumes accommodate more species. However, this pattern is consistent with niche partitioning accompanied by allee effects and/or competition^17, 22^ driving random extinction processes that lead to a reduction in clade-niche-size as postulated in Fig. 3b. In fact the significant direct effect of competition (PCI, −0.37) and relatively strong effect of clade age (CA, −0.26) on clade nice size (Fig. 3a), are consistent with such competition driven extinction processes unfolding through time^17^. The absence of this effect in the 42 clade analysis probably reflects the much younger average clade age (17my compared 112my), and smaller clade sizes, which mean that partitioning/extinction processes (Fig. 3b) will be less frequent and therefore more difficult to detect. This is also in line with previous work^23^ suggesting extinction played a pivotal role in the diversification of older conifer clades.

**Figure 3.**
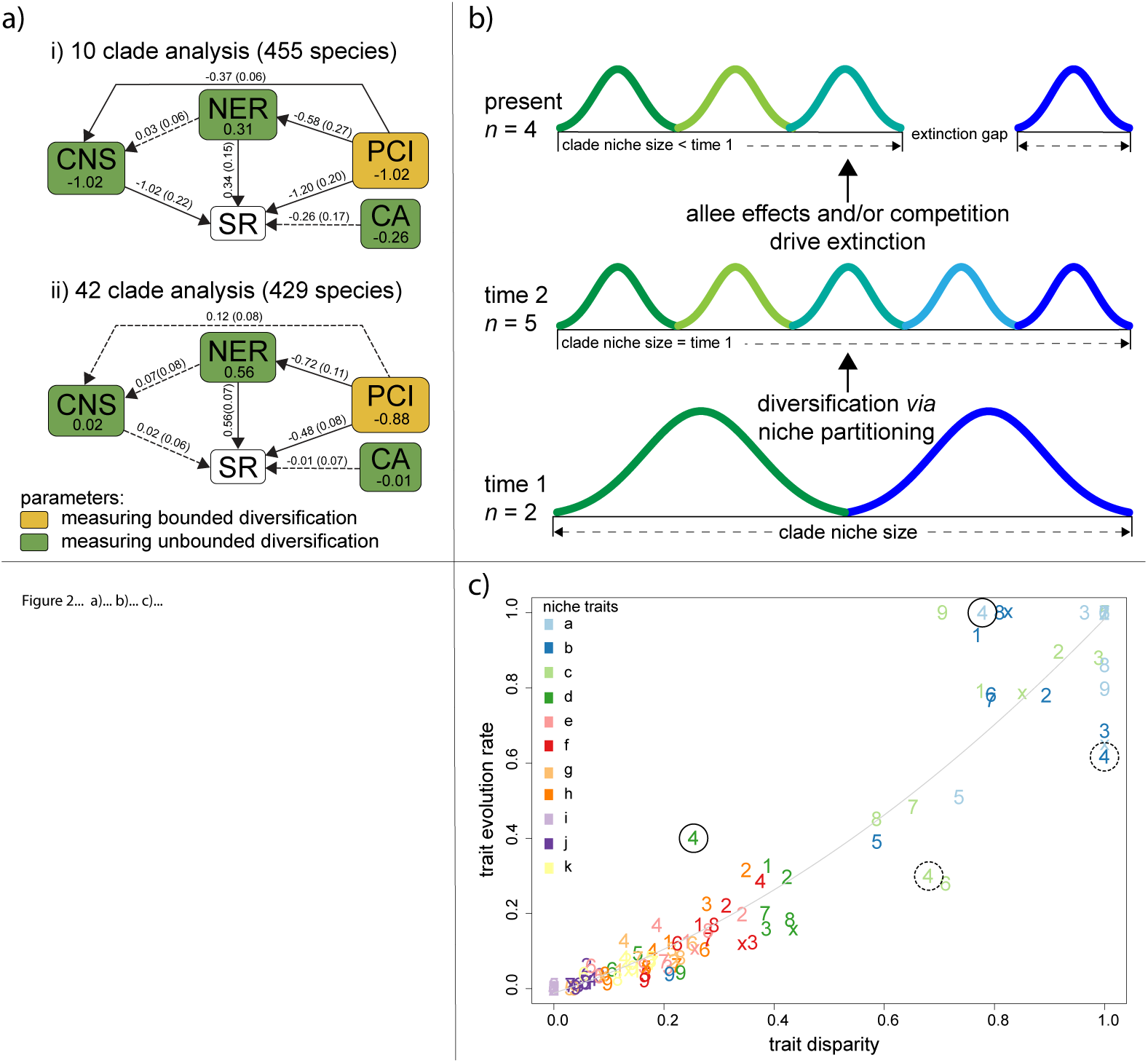
a) Bayesian path analysis showing the relative effects of niche and phylogenetic parameters on clad species richness for 455 conifer species in i) 10 large clades, and ii) 42 smaller clades. Total effect size is shown with the parameters, while direct effects and their standard deviation are shown along the vertices. Solid lines are significantly different from zero (95% credible intervals not including zero). SR = species richness; CA = clade age; PCI = phylogenetic competition index; NER = multivariate niche evolution rate; CNS = clade niche size. b) example of how niche partitioning combined with extinction associated with allee effects and/or competition, can result in a negative relationship between clade niche size and species richness. Different coloured curves represent species. c) within clade variation in the flexibility of niche traits. Coloured numbers refer to clades 1-10 (with x=10), and different colours refer to to different niche dimensions. The order and definition of niche dimensions (a-k) are as Fig 3. The best fit line (grey) is to allow visualisation of residual deviance (see text). Niche traits in solid and dashed circles are discussed in the text.

Although most previous work has favoured either bounded^2, 8, 12, 24^ *or* unbounded^3, 25, 26^ processes driving diversification, our results are consistent with previous observational^10, 18^, theoretical^14^ and modelling^4, 10^ work, which suggests that both bounded and unbounded processes influence diversification. For example, much of the empirical evidence is consistent with diversification slowing, rather than reaching an asymptote^14, 18^. This led to the “damped increase” hypothesis, which in line with our results, suggests that competition induced by niche filling reduces diversification rate, while specialisation or new ecological opportunities counteract this effect^14^. Others have extended these ideas to show that the incongruity between strict bounded and unbounded views could be overcome by allowing diversity-accumulation-models to vary between periods of either bounded or unbounded diversification^4^. These studies do not however provide a population/species level mechanism that could drive shifts in diversification processes^4^.

To address this mechanistic basis, we examined whether niche dimensionality can drive variation in diversification processes^4, 27^. To explore this possibility we examined evidence for conservatism in the evolution of the traits that define the physiological niche of species in our dataset (see Methods). We found substantial variation in the level of conservatism within and between traits (Supplementary Materials Table S1), suggesting that some niche dimensions may be more evolutionarily labile than others. Indeed, a comparison of trait disparity and trait evolution rates for each trait and clade combination (10 clade classification), showed high levels of variation in evolution rate between traits within clades (Fig. 3c). The deviations from from the trend line in Figure 3c highlights this variation. For example in Clade 4, traits a and d (highlighted with solid circles) are more labile than predicted by the degree of trait disparity, whereas traits b and d (highlighted with dashed circles) are more conservative than expected. Such variation in evolutionary flexibility between traits within clades may accommodate the operation of both bounded and unbounded processes. This can be seen more clearly by focussing attention on single clades. For example, in Clade 7 (*Pinus*, Fig. 4), the effect of soil moisture on growth (panel f) is highly conserved in the sub-clades highlighted with solid ellipses, suggesting that interspecific competition is likely to be high along this niche dimension in these sub-clades. However, these same sub-clades are labile in terms of their temperature requirements for growth (traits b and c, highlighted with dashed ellipses in Fig. 4b and Fig. 4c), indicating that evolution and specialisation are possible along these niche dimensions (Fig. 4). Analogous patterns can be seen in the other *Pinus* sub-clades (Fig. 4) and the other clades (Supplementary Materials Fig. S1-S9).

**Figure 4.**
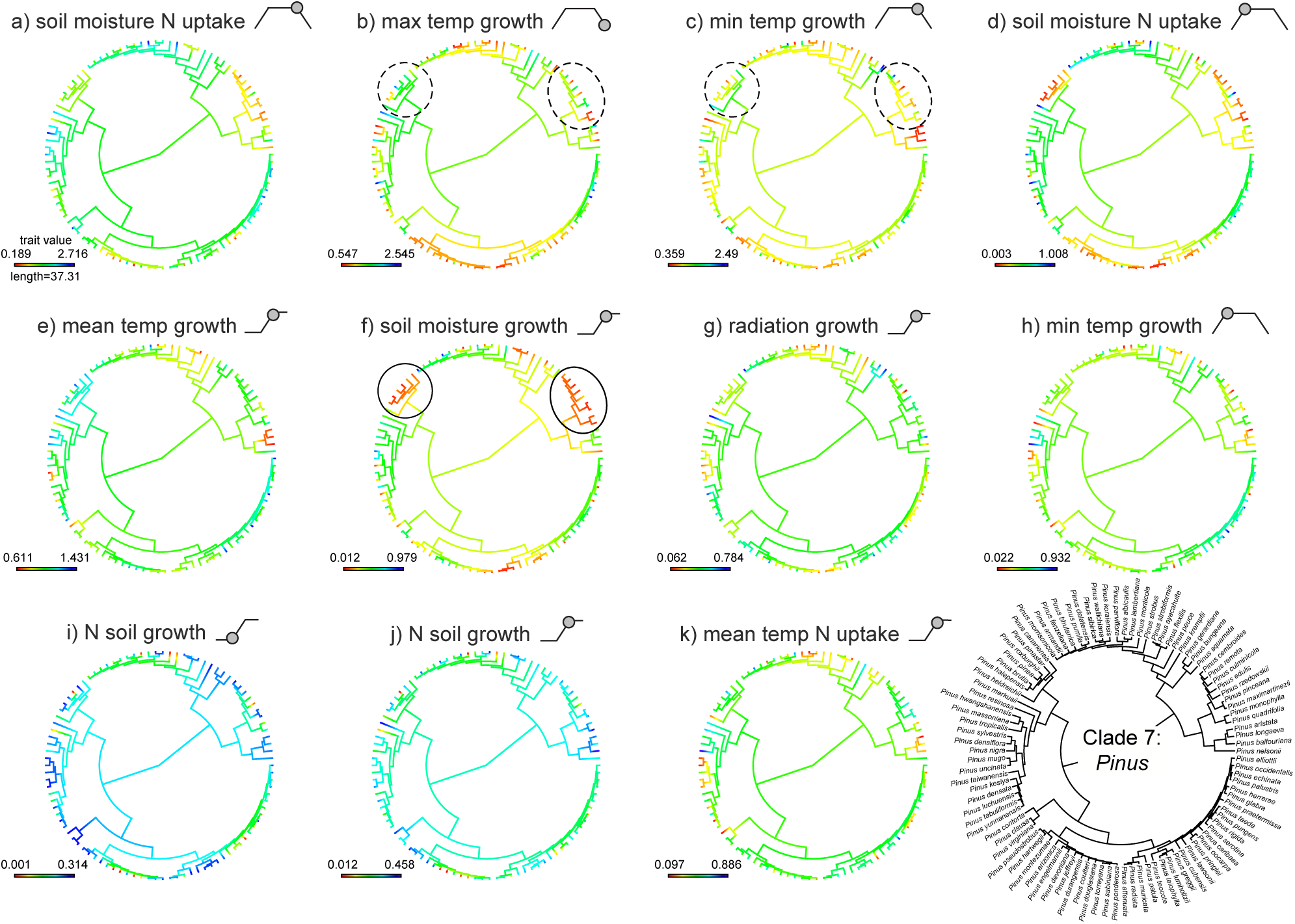
Phylogenies of Clade 7 (*Pinus*) showing ancestral state reconstructions of the 11 most important niche dimensions in order of importance (a-k). The bottom right panel shows the same phylogeny with species names. Sub clades within *Pinus* with conservative (solid ellipse) and labile (dashed ellipse) niche dimensions are highlighted and discussed in the text. The filled circle on trapezoid and logistic diagrams beside the trait names, show how the trait relates to the modelled growth or resource acquisition function. Fore example, (a) is the point at which soil moisture causes a reduction in N uptake, that is, when waterlogging reduces N uptake.

Thus, by considering the multi-dimensional nature of niche evolution, we have shown how bounded and un-bounded diversification processes may simultaneously control diversification rates. Niche dimensionality has long been thought to promote diversity by partitioning resources and facilitating coexistence^28^, and there is considerable empirical support for this hypothesis^27^. Most previous assessments of how niche characteristics impact macro-diversification have used low dimensional proxies of the niche such as body size^12^ or climatic range^11^. In contrast, our assessment of multiple, physiological niche traits, reveals that both diversity-limiting competition, and diversity-promoting evolution may operate concurrently. At the population level these processes are likely to be separated in space and/or time - in line with models by McPeek^29^ and Marshall and Quental^4^ respectively. For example, populations along environmental gradients could experience variation in the opportunity for specialisation or niche expansion along some niche dimensions but experience competition along other niche dimensions^29^. Similarly, changes in the environment could induce temporal variation in selection pressure that affects the interplay between conservative and labile niche traits^4^.

In summary, we have identified how processes that define the niche geometry of conifer clades can jointly promote and constrain diversification. Our results confirm that the contrasting processes that underpin bounded and unbounded diversification have both operated during the evolution of a major lineage. Our study thereby provides an analysis framework for a new multi-dimensional-niche hypothesis that unifies the bounded and unbounded hypotheses^4, 10, 14, 18^. The study also highlights a potential anthropogenic obstruction to future diversification. Habitat loss and fragmentation threaten existing global biodiversity by (among other things) increasing extinction rates^30^. Our results suggest that habitat loss and fragmentation could also have a compounding negative effect on future diversity accumulation. Communities that are “crowded” into smaller areas of remnant habitat are likely to experience increased competition^30^, thereby increasing the likelihood that bounded processes will constrain diversification.

## Methods

### Data acquisition and preparation

Geo-referenced collection data for all conifer species were extracted from the Global Biodiver-sity Information Facility (www.gbif.org). These data were supplemented by published species records not in GBIF from:^31–36^. Climate estimates were made for each point record, using Worldclim (37). Data was cleaned manually by firstly eliminating duplicate records, then for consistency with species distribution descriptions^31^, and then by comparing Worldclim estimates of altitude, with the altitudes provided with each site record. Where Worldclim altitudes were inconsistent with the altitude in species descriptions by more than 300 m, we replaced these records with estimates from nearby sites with altitudes consistent with the descriptions.

### Estimating physiological niche traits

We estimated the physiological niche traits of the study species using a physiologically-based approach to species distribution modelling^15^. This method uses the Thornley Transport Resistance (TTR) model of plant growth^37^ to estimate the niche traits that match the observed distribution of species. The TTR model^37^, is an ordinary differential equation model that considers how plant growth is influenced by carbon uptake, nitrogen uptake, and the allocation of carbon and nitrogen between roots and shoots. It explicitly separates the physiological processes of resource uptake from biomass growth. The implementation by^15^ relates the uptake and growth processes to environmental forcing variables. Specifically, the model considers how carbon uptake might be limited by temperature, soil moisture, solar radiation, and shoot nitrogen; nitrogen uptake might be limited by temperature, soil moisture, and soil nitrogen; and growth and respiration loss might be influenced by temperature. The model runs on a monthly time step, which allows it to explicitly consider how seasonal fluctuations in the forcing variables interactively influence plant resource uptake and growth.^15^ provides a full description of the model and its assumptions.

We use the cleaned presence dataset described above to identify locations where the species occur. A variety of methods for simulating absence points (often called pseudoabsence points) are available, but the method adopted is regarded as a relatively small source of error^38^. Our method balances the number of presence and absence points and stratifies by environment type the selection of absence points. To define environment types we use a partitioning algorithm *clara*^39^ to classify the TTR input variables into 25 environmental zones. We further restricted the selection of absence points to the zoological realm(s) where the species occurs and to distances >0.25 km from the presence points.

The model predicts the potential biomass of an individual plant as a function of the environmental forcing variables at a site. Following the work of^15^, we assume that *p_i_*, the probability of a species occurring at site *i*, is described by the complementary log–log of the modelled plant biomass at site *i* and that the likelihood of observing the presence absence data (*y_i_*) at site *i* is described by the Bernoulli distribution. To estimate the parameters, we used the differential evolution optimization algorithm^40^ to find the set of parameters that maximizes this likelihood over all sites. The model fits were evaluated by examining the confusion matrix (a matrix comparing the number of true positives, true negative, false positives and false negatives), with particular weight given to the false negative rate, i.e. instances where the model predicts the species to be absent, but it is actually present (Supplementary Materials Table S2).

We restrict projection of potential species ranges to the subset of environmental zones (see above) present in each species’ occurrence data; this prevents predictions beyond the data domain used for estimating the model parameters. We calculated the niche size of species in two ways: 1) projecting species ranges for the world, and 2) using a resampled dataset that assumes that the world’s environmental zones are equally common. This second method corrects for any bias in projected range size introduced by variation in the extent of different environmental zones, but maintains the covariance structure of the environmental data^41^. To create a dataset where each environmental zone is equally common, we created a resampled dataset of the environmental data. We again use *clara* to classify the global TTR input data into 50 environmental zones. We then sampled a finite number (1,000 in our case) of locations from each of 50 environmental zones, which produces an environmental dataset where each environment zone is equally represented. We then project the range sizes of species in this resampled environmental space. Analyses conducted using geographic locations and resampled locations yielded very similar results. The analysis based on resampled locations is presented in the main manuscript while the analysis based on geographic locations is available in Supplementary Materials Fig. S10.

### Phylogenetic methods

We used the fossil calibrated conifer phylogeny of Leslie et al.^21^, which is based on two chloroplast genes and two nuclear genes. We pruned this 487 species tree to match the 455 species for which we had good distributional data. Although a clade is any monophyletic group in a phylogeny, the ability to detect effects in clade-wise analysis will be in part reliant on having enough variation in clade size^42^. Therefore we developed two clade classifications. The first inclusive division is based on tree topology at deeper well supported nodes, and it aimed to retain major taxonomic groups such as *Pinus*, resulting in 10 clades (supplementary Information 3). The second lower division is based on a time-slice approach at Eocene/Oligocene boundary (33.9ma) because using tree topology closer to the tips becomes more difficult. This second approach produced 68 clades, 28 of which included a single species. These single species were dropped from the analysis, leaving 42 clades and 429 species in the second analysis (Supplementary Materials Table S2). We recognize that removing single species clades might bias rate estimates because these are the clades with the lowest diversification. However, the dataset still covers a wide range of clade species richness (2 - 45 species), and meaningful estimates for single species cannot be calculated for most subsequent metrics used in our analysis (e.g. niche evolution rate, clade niche overlap, clade geographic overlap etc.). Furthermore, this potential bias only affects the 42 clade analysis and the general agreement between the 10 and 42 clade analyses (see main text) suggests that any effect is inconsequential.

### Clade level data

For each clade we calculate the following metrics: age, niche size corrected for species number, niche evolution rate, and phylogenetic competition index. The crown age of the clade was calculated directly from the tree using the branching time function in APE^43, 44^. Because clades with more species could have larger clade niche sizes by chance, we used the mean of 10,000 bootstrap resamples to obtain corrected estimates of the clade niche size relative to the smallest clade (12 and 2 for the 10 and 42 clade datasets respectively).

The calculation of niche evolution rate involves using a multi-variate model. The TTR species distribution model estimates 24 parameters associated with plant growth (see above). For this reason we first extracted the most informative of the 24 niche parameters for the analysis, Specifically we used phylogenetically corrected principal components analysis (PCA) to identify which model parameters had the most influence on shaping niche space in our dataset. PC 1 to 8 explained over 94 percent of the variation in the data set. The most influential parameters were identified based on the eigan vector loadings > 0.3, and vector plots were used to exclude correlated parameters. This procedure identified 11 parameters (Fig. 3 and 4) which were ranked in order of importance by summing the effect of each trait on each PC weighted by the proportion of the variance explained by that PC. For each clade these 11 parameters were fit together in a multivariate Brownian Motion (BM) of evolution in OUCH^45^. In the 42 clade analysis, clades with fewer than 12 species had insufficient degrees of freedom to fit the model, and parameters were dropped, (starting with the lowest ranked), until a model could be fitted. Following^11^, the trait evolution model was used to calculate the variance-covariance matrix for each clade. The diagonal elements of this matrix represent the phylogenetic rate of character evolution which were summed to provide a multivariate rate parameter for each clade - the NER^11^.

The bounded hypothesis proposes that competition plays a key role in limiting diversification. Competition is likely to be most intense between close relativities due to similar physiological requirements (or niches) wherever species co-occur^17^. To estimate competition, we produce a metric which summarises the degree of expected niche overlap and observed geographic overlap between species within clades. Geographic overlap between each species pair was estimated by producing a matrix of pairwise distance between all geo-referenced occurrence records. The average of this matrix was taken for each pair to produce a pairwise matrix of geographic distances between all species. This distance matrix was transformed to scale between 0 and 1, and the inverse was taken to provide an index of observed geographic similarity (overlap). Schoener’s index^46^ of niche overlap was estimated for each pair of species from the projected species distributions (i.e. the potential niche of the species) in SPAA^47^, and the subsequent matrix was also rescaled between 0 and 1. We then took the product of these overlap estimates to produce the phylogenetic competition index, thus the index has a potential range from 0 to 1, so that species pairs with high niche overlap and high geographic overlap have a high competition score, and those with low overlaps for one or both metrics have a low competition index. The mean was taken from clade level subsets of this matrix to produce the phylogenetic competition index for each clade. It should be noted that this index is a minimum estimate, because competition with more distantly related species is also possible^24, 48, 49^ (p. 77). Incorporating community wide competition is theoretically possible using our approach (geographic overlap*niche overlap), although it would be very data intensive and is outside the scope of this work. Additionally, despite the possible underestimate of competition, our model structure(see bellow) means that the calculation of un-associated dependencies (i.e. CA on SR; NER on SR; NER on CNS; CNS on SR, see Fig. 2) are not affected.

### Regression modelling

We developed an a-priori conceptual model (Fig. 2) to estimate the relationships between SR, CNS, NER, CA and the PCI. The unbounded model predicts that specific evolutionary characteristics, controlled by phylogenetic niche conservatism, lead to clade specific diversification rates. This has two consequences, 1) when the effect of diversification rate is factored out older clades will have more species than younger clades; and 2) positive diversification will involve niche evolution that manifests as either the expansion or partitioning of clade niche space as species accumulate. In line with these predictions our model allows: 1) CA to directly influence SR; and 2) NER to influence SR both directly, and indirectly, via its effect on CNS, with the direct relationship between CNS and SR indicating the mode of niche evolution (expansion or partitioning). Conversely, the bounded diversity model predicts that competition for limited resources places a limit on species number. It has long be recognised that competition is likely to be most intense between close relatives, because the ecological requirements of relatives are likely to be similar due to phylogenetic niche conservatism. Our estimate of PCI quantifies expected competition between species within clades. Therefore we allow PCI to directly effect SR, however, because PCI quantifies interactions between niches, it is also allowed to indirectly influence SR via NER, and CNS.

We used Bayesian path analysis to calculate the effects in the path diagram (Fig. 2), while accounting for non-independence associated with phylogenetic relationships^50^. The total effect of each model parameter on the response variable (SR) was calculated from the direct and indirect effects following^51^. All model parameters were normalised and centred to a mean of zero and constant standard deviation. Following^52^, we use relative log-transformed species richness. For each analysis (10 and 42 clade), the full phylogenetic tree was collapsed to the clade level, and the inverse of the variance covariance matrix from this clade-tree was used to explicitly correct for the phylogenetic dependencies between clades. Modelling was undertaken using JAGS^53^ running three chains for 15,000 iterations, after a burnin of 25,000, and thinning the chains to every fifth sample. Normal uniformed priors we used for the path effects and convergence was assessed using a range of diagnostics in coda^54^.

### Niche trait analysis

We investigated the role of niche dimensionality in promoting both bounded and unbounded process, by using a trait like analysis to quantify the evolution of individual niche dimensions at the level of the full phylogeny and within clades. This analysis focused on the 11 niche dimensions identified above. Phylogenetic signal across the full phylogeny was estimated using Pagel’s *λ* ^55^ with significance assessed using likelihood ratio tests in PHYTOOLS^56^. The PHYTOOLS function “contMap” was used to produce ancestral state reconstructions for each of the 11 most important niche traits. A second round of niche evolution modelling focused on estimating the evolution rate of the 11 primary niche dimensions independently for each clade in the 10 clade analysis. This was done as above except single variate BM models were fitted in OUCH rather than multi variate models. the subsequent trait evolution rate’s for each clade were rescaled between zero and one to allow comparisons across clades. In order to produce a corresponding estimate of trait disparity (the magnitude of variation in actual trait values) average pairwise distance between species in each clade was calculated from raw trait data, and rescaled between 0 and 1. These two metrics were plotted in xy space to allow visualisation of clade level variation in evolutionary flexibility (Fig. 3c). We also made clade level ancestral reconstructions of the 11 main niche dimensions for the 10 large clades to assess variation in the conservation of niche dimensions within clades (Fig. 4; Supplementary Material Fig. S1-S9).

## Data availability

The data and computer code that support the findings of this study are available from the corresponding author upon request.

## Author contributions

All authors were involved in developing the ideas. GJJ provided the distribution data and phylogeny. MJL and SIH undertook the analysis. MJL lead the writing. All authors contributed to the text. SIH and DB secured funding for the research.

## Acknowledgements

We thank Bill Lee, Richard Gill, Esther Dale and members of the Eucalypt Genetics Research Group (University of Tasmania) for key discussion about the ideas. The work was funded by Te Aparangi Marsden Fund grant UOO1411.

## Author Information

Reprints and permissions information is available at www.nature.com/reprints. The Authors declare no competing financial interests.Correspondence and requests for materials should be addressed to.

## References

1. Sepkoski, J. J. A kinetic model of phanerozoic taxonomic diversity. iii. post-paleozoic families and mass extinctions. Paleobiology 10, 246–267 (1984).

2. Rabosky, D. L. & Hurlbert, A. H. Species richness at continental scales is dominated by ecological limits. The Am. Nat. 185, 572–583 (2015).

3. Harmon, L. J. & Harrison, S. Species diversity is dynamic and unbounded at local and continental scales. The Am. Nat. 185, 584–593 (2015).

4. Marshall, C. R. & Quental, T. B. The uncertain role of diversity dependence in species diversification and the need to incorporate time-varying carrying capacities. Philos. Transactions Royal Soc. Lond. B: Biol. Sci. 371, DOI:10.1098/rstb.2015.0217 (2016). DOI:10.1098/rstb.2015.0217.

5. Sepkoski, J. J. A kinetic model of phanerozoic taxonomic diversity ii. early phanerozoic families and multiple equilibria. Paleobiology 222–251 (1979).

6. Alroy, J. The shifting balance of diversity among major marine animal groups. Sci. 329, 1191–1194 (2010).

7. Rabosky, D. L. Ecological limits and diversification rate: alternative paradigms to explain the variation in species richness among clades and regions. Ecol. Lett. 12, 735–743 (2009).

8. Rabosky, D. L. & Lovette, I. J. Density-dependent diversification in north american wood warblers. Proc. Royal Soc. Lond. B: Biol. Sci. 275, 2363–2371 (2008).

9. Derryberry, E. P. et al. Lineage diversification and morphological evolution in a large-scale continental radiation: the neotropical ovenbirds and woodcreepers (aves: Furnariidae). Evol. 65, 2973–2986 (2011).

10. Ezard, T. H. G., Aze, T., Pearson, P. N. & Purvis, A. Interplay between changing climate and species’ ecology drives macroevolutionary dynamics. Sci. 332, 349–351 (2011).

11. Title, P. O. & Burns, K. J. Rates of climatic niche evolution are correlated with species richness in a large and ecologically diverse radiation of songbirds. Ecol. Lett. 18, 433–440 (2015).

12. Price, T. D. et al. Niche filling slows the diversification of himalayan songbirds. Nat. 509, 222–225 (2014).

13. Price, T. The debate on determinants of species richness. The Am. Nat. 185, 571–571 (2015).

14. Cornell, H. V. Is regional species diversity bounded or unbounded? Biol. Rev. 88, 140–165 (2013).

15. Higgins, S. I. et al. A physiological analogy of the niche for projecting the potential distribution of plants. J. Biogeogr. 39, 2132–2145 (2012).

16. Evans, M. E., Merow, C., Record, S., McMahon, S. M. & Enquist, B. J. Towards process-based range modeling of many species. Trends Ecol. & Evol. 31, 860–871 (2016).

17. Rabosky, D. L. Diversity-dependence, ecological speciation, and the role of competition in macroevolution. Annu. Rev. Ecol. Evol. Syst. 44, 481–502 (2013).

18. Valentine, J. W. Niche diversity and niche size patterns in marine fossils. J. Paleontol. 43, 905–915 (1969).

19. Wiens, J. J. The niche, biogeography and species interactions. Philos. Transactions Royal Soc. Lond. B: Biol. Sci. 366, 2336–2350 (2011).

20. Farjon, A. & Filer, D. An atlas of the world’s conifers: an analysis of their distribution, biogeography, diversity and conservation status (Brill, Leiden, 2013).

21. Leslie, A. B. et al. Hemisphere-scale differences in conifer evolutionary dynamics. Proc. Natl. Acad. Sci. 109, 16217–16221 (2012).

22. Rosenblum, E. B. et al. Goldilocks meets santa rosalia: An ephemeral speciation model explains patterns of diversification across time scales. Evol. Biol. 39, 255–261 (2012).

23. Crisp, M. D. & Cook, L. G. Cenozoic extinctions account for the low diversity of extant gymnosperms compared with angiosperms. New Phytol. 192, 997–1009 (2011).

24. Silvestro, D., Antonelli, A., Salamin, N. & Quental, T. B. The role of clade competition in the diversification of north american canids. Proc. Natl. Acad. Sci. 112, 8684–8689 (2015).

25. Belmaker, J. & Jetz, W. Relative roles of ecological and energetic constraints, diversification rates and region history on global species richness gradients. Ecol. Lett. 18, 563–571 (2015).

26. Hedges, S. B., Marin, J., Suleski, M., Paymer, M. & Kumar, S. Tree of life reveals clock-like speciation and diversification. Mol. Biol. Evol. 32, 835–845 (2015).

27. Harpole, W. S. et al. Addition of multiple limiting resources reduces grassland diversity. Nat. 537, 93–96 (2016).

28. Hutchinson, G. E. Cold spring harbor symposium on quantitative biology. Concluding remarks 22, 415–427 (1957).

29. McPeek, M. A. The ecological dynamics of clade diversification and community assembly. The Am. Nat. 172, 270–284 (2008).

30. Fahrig, L. Effects of habitat fragmentation on biodiversity. Annu. review ecology, evolution, systematics 34, 487–515 (2003).

31. Farjon, A. A Handbook of the World’s Conifers (2 vols.), vol. 1 (Brill, 2010).

32. Zhao, X. et al. Genetic variation and selection of introduced provenances of siberian pine (pinus sibirica) in frigid regions of the greater xing’an range, northeast china. J. forestry research 25, 549–556 (2014).

33. Petrova, E., Goroshkevich, S., Belokon, M., Belokon, Y. S. & Politov, D. Distribution of the genetic diversity of the siberian stone pine, pinus sibirica du tour, along the latitudinal and longitudinal profiles. Russ. J. Genet. 50, 467–482 (2014).

34. Timoshok, E., Timoshok, E. & Skorokhodov, S. Ecology of siberian stone pine (pinus sibirica du tour) and siberian larch (larix sibirica ledeb.) in the altai mountain glacial basins. Russ. J. Ecol. 45, 194–200 (2014).

35. Larionova, A. Y., Ekart, A. K. & Kravchenko, A. N. Genetic diversity and population structure of siberian fir (abies sibirica ledeb.) in middle siberia, russia. Eurasian J. For. Res. Univ. (Japan) 10, 185–192 (2007).

36. Hantemirova, E., Berkutenko, A. & Semerikov, V. Systematics and gene geography of juniperus communis l. inferred from isoenzyme data. Russ. J. Genet. 48, 920–926 (2012).

37. Thornley, J. H. Modelling shoot : Root relations: the only way forward? Annals Bot. 81, 165–171 (1998).

38. Barbet-Massin, M., Jiguet, F., Albert, C. H. & Thuiller, W. Selecting pseudo-absences for species distribution models: how, where and how many? Methods Ecol. Evol. 3, 327–338 (2012).

39. Kaufman, L. & Rousseeuw, P. J. Finding groups in data: an introduction to cluster analysis (John Wiley & Sons, 2009).

40. Ardia, D., Boudt, K., Carl, P., Mullen, K. M. & Peterson, B. G. Differential evolution with deoptim. The R J. 3, 27–34 (2011).

41. Higgins, S. I. & Richardson, D. M. Invasive plants have broader physiological niches. Proc. Natl. Acad. Sci. 111, 10610–10614 (2014).

42. Wiens, J. J. The causes of species richness patterns across space, time, and clades and the role of “ecological limits”. The Q. Rev. Biol. 86, 75–96 (2011).

43. Paradis, E., Claude, J. & Strimmer, K. Ape: analyses of phylogenetics and evolution in r language. Bioinforma. 20, 289–290 (2004).

44. Paradis, E. Analysis of Phylogenetics and Evolution with R (Springer Science & Business Media, 2011).

45. Butler, M. A., King, A. A. & Crespi, B. J. Phylogenetic comparative analysis: a modeling approach for adaptive evolution. The Am. Nat. 164, 683–695 (2004).

46. Schoener, T. W. Nonsynchronous spatial overlap of lizards in patchy habitats. Ecol. 51, 408–418 (1970).

47. Gotelli, N. J. Null model analysis of species co-occurrence patterns. Ecol. 81, 2606–2621 (2000).

48. Darwin, C. On the origin of species by means of natural selection, or the Preservation of favoured races in the struggle for life (John Murray, 1959).

49. Diamond, J. M. Niche shifts and the rediscovery of interspecific competition: Why did field biologists so long overlook the widespread evidence for interspecific competition that had already impressed darwin? Am. Sci. 66, 322–331 (1978).

50. Gonzalez-Voyer, A. & von Hardenberg, A. An Introduction to Phylogenetic Path Analysis (Springer Berlin Heidelberg, Berlin, Heidelberg, 2014).

51. Schumacker, R. E. & Lomax, R. G. A beginner’s guide to structural equation modeling (Psychology Press, 2004).

52. Rabosky, D. L., Slater, G. J. & Alfaro, M. E. Clade age and species richness are decoupled across the eukaryotic tree of life. PLoS Biol 10, DOI.org/10.1371/journal.pbio.1001381 (2012).

53. Plummer, M. et al. Jags: A program for analysis of bayesian graphical models using gibbs sampling. In Proceedings of the 3rd international workshop on distributed statistical computing, vol. 124, 125 (Vienna, 2003).

54. Plummer, M., Best, N., Cowles, K. & Vines, K. Coda: Convergence diagnosis and output analysis for mcmc. R News 6, 7–11 (2006).

55. Pagel, M. Inferring the historical patterns of biological evolution. Nat. 401, 877–884 (1999).

56. Revell, L. J. phytools: an r package for phylogenetic comparative biology (and other things). Methods Ecol. Evol. 3, 217–223 (2012).

